# A Linkable, Polycarbonate Gut Microbiome-Distal Tumor Chip Platform for Interrogating Cancer Promoting Mechanisms

**DOI:** 10.1101/2023.08.31.555803

**Authors:** Danielle S.K. Brasino, Sean D. Speese, Kevin Schilling, Carolyn E. Schutt, Michelle C. Barton

## Abstract

Gut microbiome composition has been tied to diseases ranging from arthritis to cancer to depression. However, mechanisms of action are poorly understood, limiting development of relevant therapeutics. Organ-on-chip platforms, which model minimal functional units of tissues and can tightly control communication between them, are ideal platforms to study these relationships. Many gut microbiome models have been published to date but devices are typically fabricated using oxygen permeable PDMS, requiring interventions to support anaerobic bacteria. To address this challenge, a novel platform was developed where the chips were fabricated entirely from gas-impermeable polycarbonate without tapes or gaskets. These chips replicated polarized villus-like structures of the native tissue. Further, they enabled co-cultures of commensal anaerobic bacteria *Blautia coccoides* on the surface of gut epithelia for two days within a standard incubator. Another complication of PDMS devices is high ad-/absorption, limiting applications in high-resolution microscopy and biomolecule interaction studies. For future communication studies between gut microbiota and distal tumors, an additional polycarbonate chip design was developed to support hydrogel-embedded tissue culture. These chips enable high-resolution microscopy with all relevant processing done on-chip. Designed for *facile* linking, this platform will make a variety of mechanistic studies possible.

## Introduction

Gut microbiomes have received a dramatic increase in attention for their substantial roles in maintaining human health^[1,2]^. Further, diseases ranging from atherosclerosis^[3]^ to Alzheimer’s^[4]^ to cancer^[5–7]^ have been correlated with changes in gut microbiome composition. These findings can be credited, in large part, to the vast reductions in sequencing costs over recent years. However, mechanisms of action behind these relationships are still poorly understood, leaving unexplored opportunities towards the improvement of targeted healthcare.

While *in vitro* and *in vivo* models have enabled the testing of some hypotheses behind gut microbiome-disease correlations, critical issues in their implementation remain. *In vivo* models facilitate the study of specific microbial compositions through inoculation of gnotobiotic mice to study host-microbe interactions^[8]^. However, these systems can suffer from poor translation to patient outcomes due to key differences in diet and physiology^[9]^. Alternatively, *in vitro* culture can leverage human cells. But these studies also suffer from limitations due to oversimplification using single cell types, culture conditions which differ greatly from native biology, and rapid overgrowth during co-culture studies^[10]^.

Organ-on-chip models offer a unique solution toward the study of host-microbe interactions. These platforms aim to bridge the gap between two-dimensional cell culture and patient outcomes by using human-derived cellular material, three-dimensional growth environments, and mechanical stimuli for the construction of minimal functional units of target tissues^[11]^. With the capacity to connect flow between two or more microphysiological systems (MPS)^[12,13]^, as they are sometimes called, it becomes possible to isolate particular inter-organ relationships^[14–16]^. This makes these platforms an optimal model to study mechanisms behind the association between gut microbes and disease arising in non-adjacent tissue.

Previous work demonstrated the formation of microbe-epithelial interfaces for modelling the gut microbiome^[17–21]^. These systems utilize flow to drive formation of polarized villus-like structures with strong barrier function over just a few days. Flow also enables extended co-culture with either predefined bacterial species^[17,19–21]^ or an inoculum derived from fecal samples to replicate a full microbial community^[18]^. However, a majority of devices are fabricated from polydimethylsiloxane (PDMS), which has high oxygen permeability, requiring that the system is used within anaerobic chambers or under constant flow of deoxygenated media to maintain the viability of gut bacteria, most of which are sensitive to oxygen^[22]^. Further, PDMS suffers from high levels of absorption and adsorption of molecules into the material and onto the surface, respectively^[23–25]^, limiting the application to on-chip imaging and small molecule treatment studies. To circumvent these issues some labs have developed chips from gas-impermeable materials such as polycarbonate^[20]^, however assembly involves the use of silicone gaskets or double-sided tapes, which suffer from similar or worse absorption characteristics^[26,27]^.

Much work has also been done in the field of organs-on-chip to model disease using spheroids^[28]^, organoids^[29,30]^, and micro-dissected biopsy tissue^[31]^, including extensive development of hydrogel-embedded culture in order to engineer parenchymal tissue models^[32]^. However, again, PDMS and tapes are primary components of many such devices, limiting their application in treatment studies due to issues in ab-/adsorption. Geometric constraints of complex constructs on-chip can also limit imaging applications requiring thoughtful design^[30]^.

Herein, we report the development and application of MPS devices composed solely of polycarbonate. This material facilitates control over oxygen exposure to culture gut microbiome interface models using standard tissue culture infrastructure. Further, tumor chip devices support culture of a variety of hydrogel embedded tissue models. Polycarbonate’s material properties optimize the system for future applications in multi-organ studies to understand mechanisms behind gut dysbiosis and cancer progression (Figure 1).

**Figure 1.**
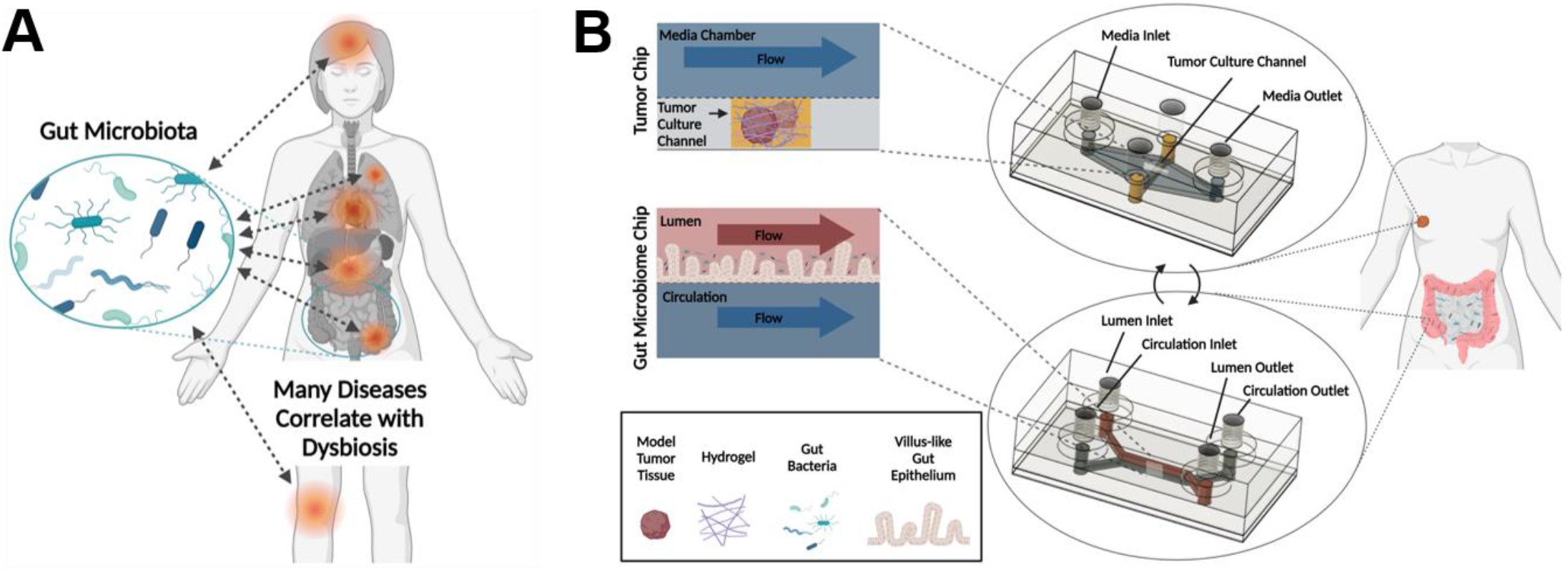
Overview of gut-microbiome disease interactions and a platform to study mechanisms behind these relationships. A) Illustration of the variety of diseases associated with the gut microbiome including depression, Alzheimer’s, cancer, atherosclerosis, diabetes, obesity, IBD, arthritis and more; B) Illustration of the tumor chip and gut microbiome chip platforms presented herein, designed for studies into gut microbiome-tumor associations.

## Results and Discussion

### Fabrication of novel, all-polycarbonate chip platforms

Most MPS platforms leverage soft lithography techniques to generate chips from PDMS due to the rapid production capabilities of this method. However, this comes with the cost of high ad-/absorption characteristics, affecting the transport and delivery of molecules based upon their hydrophobicity^[33]^. Additionally, PDMS is highly oxygen permeable, requiring added infrastructure or methodologies to culture anaerobic species^[18,21]^. To circumvent these issues, novel chips were developed entirely from a thermoplastic polymer, polycarbonate. While others have leveraged thermoplastics in this regard, plastics were interfaced using double sided tapes^[26]^ or channels were introduced using silicone rubber gaskets^[20]^. These materials also suffer from high absorption characteristics^[27]^ and introduce multiple material interfaces for cell growth which could have negative implications when trying to culture a continuous epithelial or endothelial tissue.

In order to generate a fully polycarbonate device, all channel components were modeled using Autodesk manufacturing software, Fusion 360, followed by precision milling using a computer numerical control (CNC) micro-milling machine. This process supports the rapid design and prototyping of new chip geometries. Further, this process eliminated use of hazardous piranha or hydrofluoric acid etching agents critical to photolithography.

To generate fully polycarbonate devices, it is necessary to bond the independent components. Common methods for thermoplastic bonding include heating of the component parts to the plastic’s melting point or application of relevant solvents to the part surfaces prior to joining. However, these methodologies are unsatisfactory in the generation of microfluidics as they can severely impact the channel morphology. Vapors of appropriate solvents can swell a thin layer of the thermoplastic surface. Exposure to solvent vapor followed by compression at a temperature far below the melting temperature bonds parts without impacting channel geometries^[34,35]^. This technique was applied to the chip system in two steps to minimize warping of the thin, porous membrane which was sourced commercially from Sterlitech. A thick polycarbonate slab was threaded for tubing interfaces and bonded to the microfluidic chips using traditional solvent bonding for durability during hand-tightened tubing installation (Figure 2A). Taken together, this multi-step fabrication process enabled the fabrication of gut microbiome (Figure 2B) and tumor chips (Figure 2C,D) made entirely of polycarbonate.

**Figure 2.**
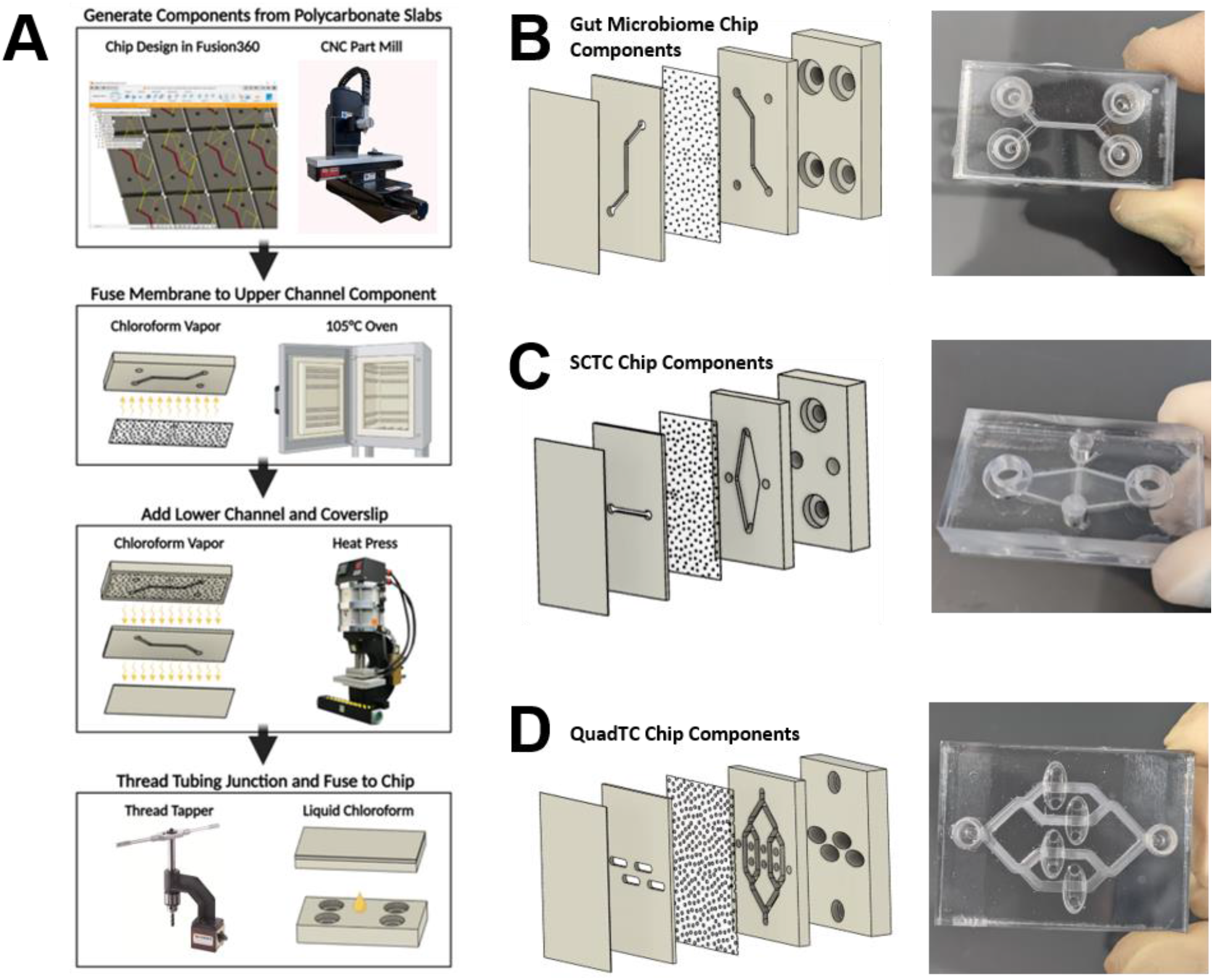
Fabrication of polycarbonate devices using machine milling and chloroform-assisted bonding. A) Manu-facturing process of polycarbonate chips. Polycarbonate slabs are milled using a Computerized Numerical Control (CNC) milling machine and tubing interfaces are adapted for fittings using a manual thread tapper. Components are assembled using vapor or liquid chloroform deposition followed by application of heat and/or pressure. CNC, heat press and thread tapper images are reproduced from Minitech Machinery Corp., Rosin Technologies LLC. and Vertex Machinery Work Co., respectively; B) Gut chip layers and fully assembled gut chip; C) Single channel tumor chip (SCTC) layers and assembled SCTC chip; D) Quad channel tumor chip (QuadTC) layers and assembled QuadTC.

### Polycarbonate gut microbiome MPS model

Gut microbiome model devices described herein use the stacked channel architecture developed by previous epithelial models (Figure 1B). These upper and lower chambers replicate the gut lumen and adjacent capillaries, respectively. Following the transition to all-polycarbonate interfaces, it was necessary to evaluate the extracellular matrix deposition process critical to facilitate cell attachment to the porous membrane surface. Assessment using cell seeding assays and two-photon microscopy indicated best membrane functionalization when pure collagen was deposited at room temperature in phosphate buffered saline (PBS) overnight, evidenced by cellular monolayer coverage and collagen fibril deposition, respectively, (Figure S1, Supporting Information).

Following channel functionalization, cells from the colon epithelial cell line CaCo-2 were cultured in the upper, luminal compartment of the gut chip (Figure 3A). Constant flow of culture media at a rate to match shear strain experienced in the colon^[36]^ produced villus-like structures (Figure 3B) over the course of 3-6 days. To characterize polarization of cells within these constructs, immunofluorescence imaging was employed. By the seventh day of culture markers of polarization were evident. Staining at basolateral interfaces by anti-beta catenin indicated that the apical surface was appropriately directed toward the upper, gut lumen compartment (Figure 3C). Additionally, staining for marker zonula occludens-1 (ZO-1) indicated the formation of tight junctions at the intercellular interfaces, critical to the proper function of barrier tissues like the gut epithelium, expressed particularly at the interface of the lumen and closer to the apical end of the cell than the nucleus, indicative of polarization (Figure 3D). Beyond morphological features of the gut model, an essential component of the microbial niche is the extracellular mucin layer produced by specialized goblet cells. Transdifferentiation of Caco-2 cells into a goblet cell-like phenotype is indicated by immunofluorescence staining of the protein mucin 2. Staining of intracellular granule structures, evident as bright spots above background (staining highlighted by arrows), within this subset of cells indicated pre-secreted mucin (Figure 3E).

**Figure 3.**
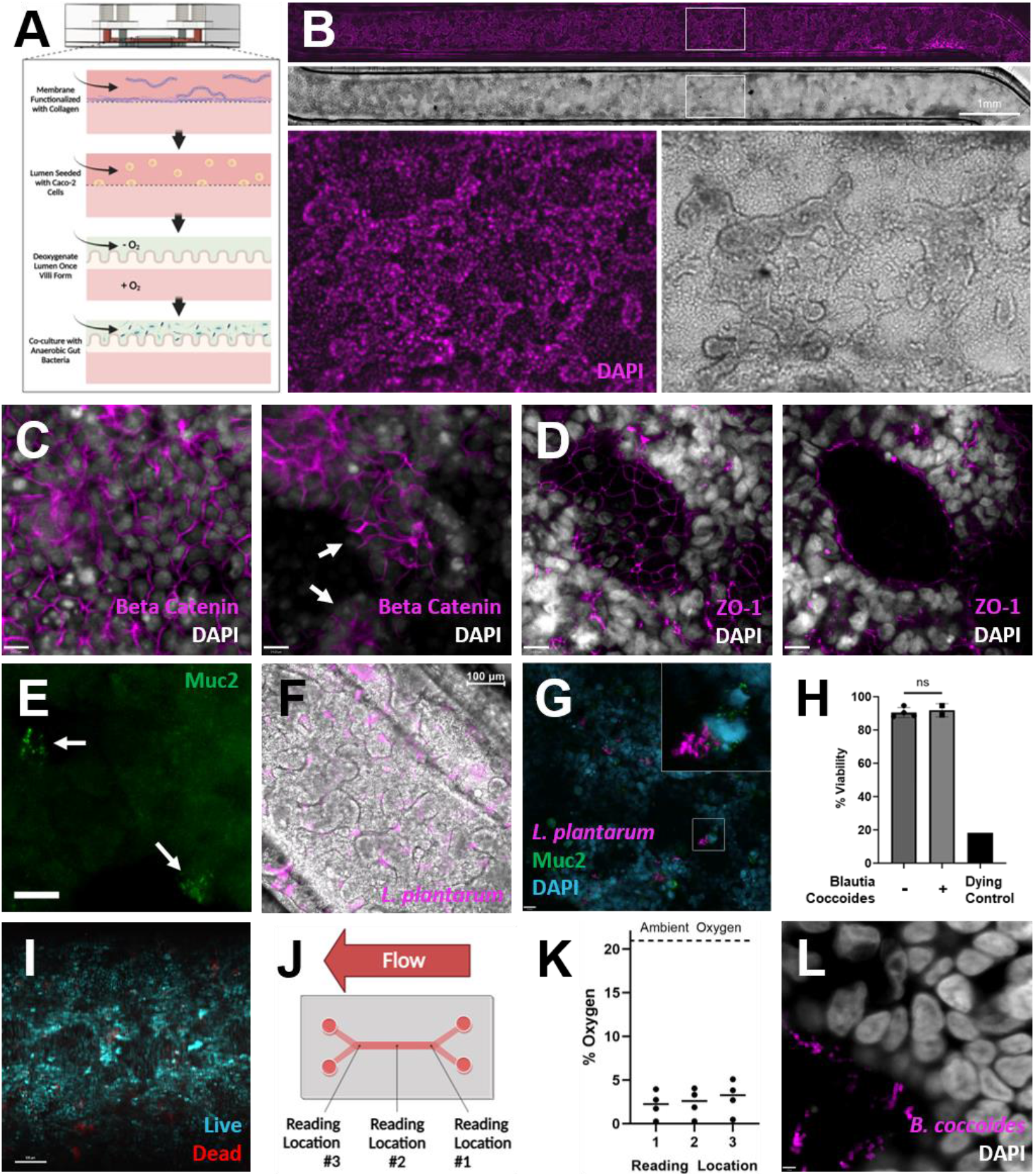
Characterization of gut microbiome chips co-cultured with facultative and obligate anaerobic bacteria. A) Schematic overview of gut microbiome assembly on chip. Channels are functionalized with collagen prior to Caco-2 cell seeding. Following growth of villus-like structures, gut lumen channel media is depleted of oxygen for co-culture with anaerobic gut microbes; B) Gut chip channel with Caco-2 cells grown into villus-like structures, shown in brightfield and with nuclei stained by DAPI (magenta), scale bar denotes 1mm; C) Immunofluorescence imaging of polarized Caco-2 layers on day 7 using markers DAPI (white) and anti-Beta Catenin (magenta) show basolateral staining of beta catenin at intercellular junctions (left) and no staining at apical surfaces (right). Arrows indicate examples of apical surfaces devoid of beta catenin staining, scale bars denote 14µm; D) Caco-2 barriers characterized by immunofluorescence imaging of tight junction protein ZO-1 (magenta) and DAPI (white), shown here on day 9, show ZO-1 expression at cellular interfaces (left) and apical to nuclei at the villus surface (right), scale bars denote 14µm; E) A subset of Caco-2 cells transdifferentiate into goblet-like cells producing mucin, noted by arrows, shown here on day 9, scale bar denotes 14µm; F) Overlaid brightfield and immunofluorescence image of Caco-2 villi co-cultured with mCherry expressing *Lactobacillus plantarum*, scale bar denotes 100 µm; G) Immunofluorescence image of Caco-2 cell nuclei (cyan), *Lactobacillus plantarum* (magenta), and mucin granules produced by goblet-like cells (green), scale bar denotes 21 µm; H) Gut chip viability assessment validates low cell death upon exposure to luminal oxygen depletion and no reduction in viability in co-culture with commensal anaerobe *Blautia coccoides*, one-way ANOVA with a Bonferroni post hoc, n=2-4 chips; I) Representative image of 9-day old gut chips cultured with low luminal oxygen showing live (blue) and dead cells (red), scale bar denotes 100µm; J) Schematic depicting oxygen reading locations taken along the overlapping region within the gut chip; K) Oxygen readings taken within the lumen of a 9-day old gut chip show an average 2-3% oxygen along the length of the chip; L) Immunofluorescence image of *B. coccoides* cocultured within a gut chip for two days. Bacteria integrated FITC-d-alanine (magenta) into cell walls during the second day of culture and Caco-2 nuclei were stained with DAPI (white), scale bar denotes 3.5µm.

Following growth of the villus-like gut epithelium, a microbial community may be introduced to the gut lumen to simulate the gut microbiome. For any bacteria of interest tolerant to oxygen, such as the commensal bacteria *Lactobacillus plantarum*, chips may be inoculated immediately following villus growth. By utilizing a strain genetically engineered to express the fluorescent protein mCherry, bacteria could be tracked using fluorescence microscopy. Bacteria were found to reside within inter-villus spaces (Figure 3F). Commensal bacteria, including several *Lactobacillus* species, have been shown to enhance expression of gut-protective mucins^[37,38]^. Immunofluorescent imaging of co-cultures within our platform show large numbers of mucin granules within goblet-like cells alongside mCherry expressing *L. plantarum* (Figure 3G).

The vast majority of gut bacteria are obligate anaerobes, many intolerant to any handling done at the bench. To culture these bacteria, all handling must be done under an inert atmosphere devoid of oxygen. Gut epithelia, on the contrary, require oxygen. To replicate the gut microbiome interface, it became necessary to provide an oxygen-free environment for the bacterial compartment, while delivering oxygen to any mammalian cell culture in a simulation of the enteric circulatory system. By utilizing oxygen impermeable materials, our platform enables maintenance of an anaerobic condition following removal of oxygen from the media reservoir. To simulate the gut microbiome interface, media reservoirs for the lumen compartment were depleted of oxygen following growth of gut villi, while lower channel media supplied oxygen to Caco-2 cells (Figure 3A). Assessments following oxygen depletion show Caco-2 cell layers are over 80% viable (Figure 3H,I). Using OXNANO probes dispersed in the media supplied to the lumen compartment, an optical sensor detected bulk oxygen concentrations of 2-3% on average, which is within physiological ranges^[39]^ (Figure 3 J,K). Under the microaerophilic conditions of the colon, diverse microbial communities depend upon a range of oxygen microenvironments^[40]^. Using phosphorescence lifetime imaging to assay oxygen tension across the cell surface, oxygen gradients could be discerned demonstrating future capabilities of the chip to support full microbial consortia (Figure S5, Supporting Information).

To demonstrate capability of co-culture with obligate anaerobes, gut lumens were inoculated with the commensal bacteria, *Blautia coccoides*. Following two days of co-culture, media was collected from each chamber and plated under anaerobic conditions to assay presence of viable bacteria. Effluent from the lumen compartment grew substantial colonies while effluent from the lower chamber did not (Figure S6, Supporting Information). Epithelia showed no reduction in viability in the presence of bacteria (Figure 3H). These results successfully demonstrate longitudinal co-culture of two biological components with discrete culture requirements to recapitulate the gut microbiome interface. To assay metabolic activity of the bacteria on-chip, fluorescently conjugated D-alanine was added to the culture media after a day of co-culture. Following a day of culture in the presence of fluorescent D-alanine, microscopy indicates substantial uptake of the molecule and integration into the bacterial cell wall. This enables on-chip tracking of the bacteria, which reside at the villus surface and in the intervillous space, and demonstrates their continued metabolic activity while on-chip (Figure 3L and Video V1, Supporting Information).

### Polycarbonate tumor chip model for hydrogel-embedded culture

*In vitro* systems, MPS included, involve a reduction in complexity to model tissues of interest. To target mechanisms of interest or optimize outcomes for patients using personalized medicine, the degree to which complexity is reduced varies. Based upon experimental constraints, different tissue models are employed ranging from cell line-derived spheroids to patient-derived tissue. To optimize utility of our device in future applications leveraging these varied model tissues, our system incorporates hydrogel-embedded culture with nutrient and waste flux via unidirectional fluid flow. The high degree of absorption and adsorption inherent in materials like PDMS^[23–25]^ commonly used for culture platforms limits applications in drug dosing studies and analytical techniques such as imaging. By constructing platforms using our polycarbonate chip fabrication protocol, we optimized our system for a wide variety of applications by reducing ab-/adsorption issues (Figure 4B). Specifically, engineering the platform for high-resolution imaging on-chip makes it possible to conduct long-term live-cell imaging to study characteristics at a range of timescales. Using cancer cell line-derived spheroids or patient-derived organoids, this platform will facilitate studies into the relationship of gut microbiome composition and distal solid tumors is organs such as the breast^[7]^.

**Figure 4.**
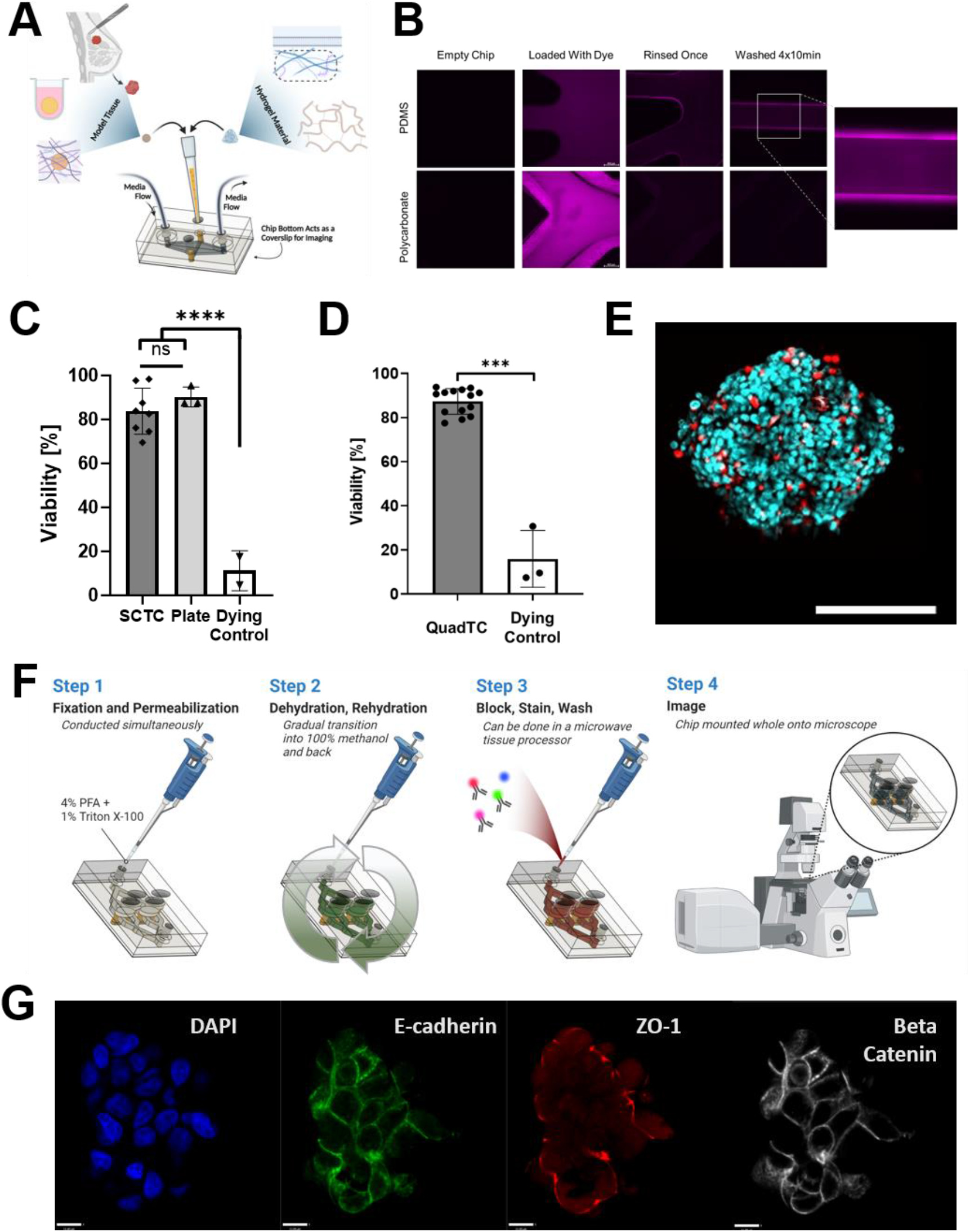
Characterization of tumor chips capable of high-resolution on-chip imaging. A) Schematic overview of tumor chip capabilities. Tissue channels may be loaded with a range of materials including spheroids and organoids, extracellular matrix proteins and synthetic polymers. Devices are designed for longitudinal studies including on-chip imaging; B) Absorption and adsorption by PDMS and polycarbonate following 2-hour dye incubation and washing by PBS. Strong residual fluorescence only on PDMS channel surfaces, scale bar denotes 400µm; C) SCTC following two days of on-chip culture in 1.45mg/mL collagen maintain equivalent viability to spheroids maintained in liquid culture, ****, p<0.0001, one-way ANOVA, taken from three chips and three plate spheroids; D) High viability of spheroids cultured in 2.7mg/mL collagen in QuadTC for two days, ***, p<0.005, t-Test; N=1-3 spheroids per well, across 4 wells; E) Representative image of spheroid cultured in QuadTC chip showing live cells (blue) and dead cells (red). Scale bar denotes 200µm; F) Schematic of on-chip staining protocol for immunofluorescent imaging; G) Confocal microscopy image of MCF7 spheroid loaded with collagen in QuadTC chip. Stained for DAPI (blue), Ecadherin (green), ZO-1 (red) and Beta-Catenin (white), scale bar represents 11µm.

Model tissues and support matrices are loaded into a recessed channel while independent ports for tubing feed a diamond shaped channel for media (Figure 4A, Figure S7 Supporting Information). Hydrogel and media compartments of the single-channel tumor chip (SCTC) are separated by a porous membrane to avoid occlusion of flow. Targeting flow perpendicular to the tissue culture channel limits accumulation of excreted factors from treated tissues. This is an issue of previous model chips that culture tissues in series as it results in heterogenous culture conditions.

By using polycarbonate it becomes possible to tightly control nutrient and oxygen flux due to the reduced permeability of polycarbonate. To ensure this reduced oxygen influx did not negatively impact tissue viability, spheroids comprised of MCF7, a breast cancer cell line, were cultured in SCTC embedded in a collagen matrix for two or three days. Staining with Hoechst 33342 and ethidium homodimer established no reduction in viability compared to spheroids maintained in liquid culture in low adhesion plates (Figure 4C). While spheroids make many mechanistic studies possible, personalized medicine requires the use of patient derived material either as whole tissue or dissociated and developed into self-assembled organoids. To test these more complex models, a patient-derived kidney cancer organoid line was cultured in basement membrane extract within the SCTC for one week. These organoids also maintained high viability (Figure S9, Supporting Information).

A second chip design, QuadTC, cultures four, independent recessed hydrogel channels, expanding upon the tumor chip capabilities by enabling replicates within a single chip (Figure 2D). Hoescht 33342 and ethidium homodimer staining confirm high viability across the channels of MCF7 cell line-derived spheroids embedded in collagen (Figure 4D,E).

Because model tissues are cultured adjacent to the coverslip bottom and polycarbonate does not suffer from high fluorophore or biomolecule adsorption, staining and high-resolution imaging can be done on-chip. Tissue processing protocols included simultaneous fixation and treatment to permeabilize membranes followed by methanol dehydration and rehydration in phosphate buffered saline. These techniques improved imaging at depth in dense spheroids. Antibody staining could be done inside a microwave tissue processor. Taken together these techniques, which can all be done on-chip due to compatibility with our chip geometry, enable acquisition of high-resolution microscopy data without removal from the chip, avoiding processing which could damage tissues or alter the microenvironment (Figure 4F). As a demonstration of function, staining for beta catenin, ZO-1, and e-cadherin showed polarity of spheroids loaded into a QuadTC (Figure 4G).

## Conclusion

This work demonstrates the capabilities of a novel, all-polycarbonate organ-on-chip platform. Using vapor-assisted fusing, milled components were assembled without the use of tapes or gaskets, minimizing ab-/adsorption issues typically found in PDMS devices. Two-channel epithelial chip models replicated villus structure growth from Caco-2 cells. With reduced oxygen permeability, polycarbonate also supported coculture of an obligate anaerobe on the surface of gut epithelia without specialized incubation conditions. A second polycarbonate chip for culture of hydrogel-embedded tumor models supported viability of cell line-derived spheroids or patient-derived organoids. Due to the flexibility of this culture system, the platform could be applied in the future to a wide range of diseases associated with gut dysbiosis, using previously developed spheroid or organoid models. In addition, similarities to biopsy culture devices suggests even wider options in selecting model tissues. Because of the material properties and culture geometry, these devices can be used for a myriad of analytical readouts with a particular focus on on-chip processing and imaging for high-resolution fluorescence microscopy. These capabilities enable long-term live-cell imaging experiments to study phenotype changes continuously instead of relying on terminal experiment analyses.

By supporting anaerobic bacteria, polycarbonate chips can culture gut microbiomes which are composed predominantly of oxygen-sensitive microbes. Single and quad-channel tumor chips can leverage the vast field of organoid development to model a diverse range of diseases by supporting hydrogel-embedded culture. By eliminating absorption and adsorption issues, biomolecules can be freely fed between these two chips using simple tubing interconnects (Figure S12, Supporting Information). Because flow is driven unidirectionally, this enables direct study of the effect one tissue has on another. Altogether, this novel chip platform will empower mechanistic studies into gut microbiome-disease correlations for improved intervention strategies and better patient outcomes.

## Materials and Methods

### Materials

Caco-2 cells were acquired from Emulate Inc. with purchase of an Emulate Bio-kit. All other materials were sourced from standard vendors including; polycarbonate (McMaster-Carr), porous membranes (Sterlitech), and *Blautia coccoides* (ATCC). *Lactobacillus plantarum* expressing mCherry were a kind gift from Dr. Michael Brasino and MCF7 cells were a kind gift from Dr. Hisham Mohammed. Organoids and organoid culture materials were a kind gift from Dr. George Thomas. Bulk oxygen tension measurements were conducted using OXNANO oxygen nanoprobes and sensor developed by PyroScience.

### Cell Culture

Mammalian cells were maintained prior to use in standard tissue culture flasks. Caco-2 cells were cultured in Gibco Dulbecco’s Modified Eagle Medium (DMEM) supplemented with 20% fetal bovine serum (FBS) and 1% penicillin/streptomycin. MCF7 cells were cultured in DMEM supplemented with 10% FBS and 1% penicillin/streptomycin. All cells were passaged at 60-80% confluence using Gibco TrypLE Express Enzyme.

### Spheroid Culture

MCF7 cells were collected once 80% confluent and counted using an automated cell counter *via* trypan blue assay. Cell suspensions were diluted using standard cell culture media to 1.2e5 cells/mL. 25uL of cell suspension was pipetted into the center of each well of Nunclon Sphera low-adhesion round-bottom 96-well plates. Following culture for three days, fresh media was added to bring the volume up to 200uL. Alternatively, at day three spheroids were washed three times with PBS followed by culture in phenol red free DMEM supplemented with 1% Gibco glutaMAX and 10% charcoal stripped FBS (white media).

### Bacteria Culture

Prior to chip inoculation, *Lactobacillus plantarum* were grown overnight in Research Products International Corp Lactobacillus De Man, Rogosa and Sharpe (MRS) broth at 37°C in a CO2 jacketed incubator. All handling for obligate anaerobe, *Blautia coccoides* was conducted in a glove bag purged with ultra-high purity nitrogen gas and oxygen was removed from culture materials using overnight exposure to AnaeroPack-Anaero in a sealed chamber. Prior to chip inoculation, *B. coccoides* were grown overnight in BD Bacto Brain Heart Infusion (BHI) broth at 37°C in an anaerobic jar. All plating for Colony Forming Unit (CFU) assays of *B. coccoides* were done on BHI agar plates, followed by growth at 37°C in an anaerobic jar overnight or for several days prior to imaging at ambient conditions.

### Chip Fabrication

Microfluidic chip parts were designed using Autodesk Fusion 360 software prior to milling on a Minitech Computer Numerical Control (CNC) micro-milling machine. Completed parts were soaked overnight in isopropyl alcohol to remove residual adhesive residue, then dried at 80°C for 30 minutes. Chloroform vapor was exposed to channel surfaces by suspending the part over a liquid chloroform bath in a sealed chamber. Chambers were allowed to develop for at least 10 minutes prior to each use. Upper gut chip channel surfaces were exposed to vapor for 5 minutes, then assembled with a hydrophilic 0.2µm pore size, polycarbonate membrane in a clamping device and baked for 12 minutes at 105°C. Channel-membrane fusions were allowed to rest overnight, then membrane was removed from lower channel port access points using a drop of dichloromethane (DCM). Lower channel pieces and bottom cover slips were exposed to chloroform vapor for 8 and 3 minutes, respectively, then assembled with channel-membrane fusions and pressed using a pneumatic heat press at 100psi and 120°C for 60 seconds. Completed chips were stored in a clamping device at room temperature. Tubing interfaces were threaded using a manual thread tapper, their surface exposed to a thin layer of liquid chloroform, then assembled with completed chips. Whole assemblies were then placed in a clamping device to dry overnight. For tissue chips, the same steps were taken with minor modifications. The initial fusion step was conducted with lower channel components and 5um polycarbonate membranes, followed by simultaneous fusion to the upper flow channel and bottom coverslip.

### Gut Chip Culture

Assembled polycarbonate gut chips were sterilized by soaking in 70% ethanol for 30 minutes. Sterilized chips were dried in a tissue culture hood for a few hours or up to overnight. To functionalize polycarbonate membranes for cell seeding, collagen (50µg/mL diluted in PBS) was introduced to each channel. Chips were placed in a humid petri dish to limit evaporation, sealed, and left at room temperature overnight. The following day, collagen solution was removed by aspiration and chips were allowed to dry in a tissue culture hood for half an hour prior to assembly with fluorinated ethylene propylene (FEP) tubing, syringes, and glass reservoirs. Media withdrawn from reservoirs into syringes was incubated in chip channels for 1 hour at 37°C. During media incubation, Caco-2 cells were collected and counted using an automated cell counter *via* trypan blue assay. Cell suspensions in standard media were prepared in new reservoirs at a concentration of 3e6 cells/mL. Following media incubation, reservoirs for each upper channel were swapped for those with cell suspension which was gently withdrawn into each chip. Cells were allowed to adhere to the channel surface for 2-2.5 hours followed by replacement of channel media with standard culture media. Flow was then applied to chips at a rate of 100uL/hr using a microfluidic syringe pump. Chips were cultured at 37°C until villi developed, typically six days. Only chips with full cell coverage were used for subsequent experiments.

### Gut Microbiome Chip Bacteria Inoculation

Gut chips were cultured for six days to develop villus-like structures at which point all reservoirs were exchanged for antibiotic free media, either DMEM supplemented with 10% FBS or white media. Before use, media for upper channels was depleted of oxygen overnight in an anaerobic jar with Aneropack-Anaero anaerobic gas generator. On the second day of culture in antibiotic free media, bacterial cultures were pelleted (4,000rcf for 5 minutes) followed by resuspension in antibiotic free cell media. Centrifugation and resuspension were repeated twice to wash the bacteria. Final bacterial suspensions were quantified *via* NanoDrop and diluted to a final O.D. of 0.5. All media exchanges were conducted in a glove bag purged with ultra-high purity nitrogen, and final diluted bacteria were prepared in a sealed reservoir. Once sealed, reservoirs were removed from the glove bag and attached in place of upper channel reservoirs for gut chips. Bacteria were gently withdrawn into the chip channel and incubated statically for 1.5hrs to allow bacteria to adhere. Unbound bacteria were subsequently washed away and flow resumed at a rate of 100uL/hr. For inoculation of *Lactobacillus plantarum* the same protocol was used, except upper channel reservoirs and bacterial inoculants were prepared in ambient conditions.

### Tissue Chip Culture

Assembled polycarbonate gut chips were sterilized by soaking in 70% ethanol for 30 minutes. Sterilized chips were dried in a tissue culture hood for a few hours or up to overnight. Once dry, chips were assembled with FEP tubing, syringes, and collection reservoirs. Spheroids were collected into a tube using a wide bore tip to minimize shearing and excess media was removed. A collagen stock solution in acetic acid was neutralized with sodium hydroxide solution and diluted with 10x PBS to achieve an isotonic solution, prior to mixing with the prepared spheroids. Collagen-spheroid preparations were allowed to warm at room temperature for three minutes prior to injection into chip culture channels to limit spheroids settling at the bottom of each channel. Chips were incubated at 37°C for one hour to allow the collagen to gel, followed by introduction of media into the upper chip channels and layered onto gel injection ports and initiation of flow at a rate of 60 or 100uL/hr.

### Two-photon Laser Scanning Microscopy

A two-photon microscope consisting of a tunable chameleon Ti:Sapphire laser (100 fs, 80 MHz, Coherent, Santa Clara, CA) on a commercial LSM 880 unit (Zeiss, Santa Clara, CA) was used for imaging. A 10X/0.45NA dry lens was used to image collagen via Second Harmonic Generation (SHG) as well as fluorescence from live and dead cells via Hoechst 33342 and Ethidium Homodimer (EthD-1), respectively. Images (1024x1024 pixels) were acquired with the laser tuned at 810 nm for SHG and 740 nm for live and dead cell imaging. Fluorescence emission from Hoechst and EthD-1 as well as SHG were set to be collected at 450/50 nm, 620/30 nm, and 405/30 nm, respectively.

### Cell Viability Determination

Chips were disassembled from tubing followed by staining with 4µM ethidium homodimer and 29µM Hoechst 33342 in culture media at 37°C. Dying chip controls were achieved by preincubation with 70% ethanol. Dying controls in gut chips and tumor chips using more dense, 2.7mg/mL collagen were additionally stained in the presence of 1% Triton X-100. Images containing stained nuclei of either live (Hoechst 33342) or dead (EthD-1) cells were processed via Volocity (Quorum Technologies, Inc., Puslinch, ON). Cells that had positive staining were thresholded, segmented, and counted with included segmentation tools in Volocity. To determine the viability, the number of live cells were divided by the sum of live and dead. These analyses were performed for each spheroid separately. Statistics were performed using either t-test or one-way ANOVA.

### Two-photon Phosphorescence Lifetime Imaging of Oxygen Sensors (2P-PLIM)

A two-photon excitable phosphorescent oxygen reporting micelle complex which contains encapsulated Ru[dpp]^2+^ was fabricated as previously described by Khan et al.^[41]^ A gut chip grown for ten days with three days of *B. coccoides* co-culture was stained with Hoechst for 30 minutes, followed by introduction of probe solution to the upper, lumen channel. Chip was disassembled from tubing and ports were sealed with cellophane tape in a gas bag to limit introduction of oxygen prior to imaging. 2P-PLIM of oxygen sensors within the chip was performed using an SPC-150 fluorescence lifetime imaging (FLIM) module (Becker & Hickl) with an added DDG-210 pulse generator on the LSM 880 microscope. Excitation of the oxygen sensors occurred at 740 nm with a 3 µs excitation gate for a total of 25.6 µs collection per pixel. Emission from the oxygen sensors was transmitted through a dichroic and further filtered via a 605/70 nm bandpass filter where the light was directed to a BiG detector (Zeiss) which was connected to the photon counting boards. To achieve adequate signal-to-noise ratios, 100 frames were collected per image. Data acquisition was performed using Becker&Hickl software (SPCM). The data was further processed via FLIMFit 5.1.1, a custom MATLAB GUI-based software tool developed by Warren et al., where the phosphorescence lifetime images were generated^[42]^.

### Immunostaining

#### Gut Chip

Caco-2 cells were fixed in-chip with 2% paraformaldehyde for 15 minutes at room temperature. Following fixation, permeabilization and blocking was performed with .2% Triton X-100 in PBS containing 1% BSA for 30 minutes. Fluorophore conjugated primary antibodies were then incubated O/N at 4°C. The following day, samples were washed 3 x 10 minutes in PBS. Included in the first wash was 1ug/ml DAPI. For high resolution imaging, a razor blade was used to carefully separate the bottom two layers of the gut chip from the rest of the chip containing the hydrophilic polycarbonate membrane that the cells were seeded on, the top channel, and the section of the chip containing the media inlet and outlet ports. It was critical not to damage the polycarbonate membrane when using the razor blade to separate the chips. The separated chip was then placed with the membrane containing the cells against a 22x40 mm #1.5 coverslip that had a drop of mounting media (Vectashield) to help hold the coverslip in place. Vectashield was then pushed through the media inlet port, ensuring it filled the entire upper channel, and the coverglass was fixed in place using nail polish to seal it to the gut chip.

#### Tumor Chip

Staining of spheroids/organoids in the SCTC and QuadTC chips was accomplished by first pushing 4% paraformaldehyde/.1% Triton X-100 into the chip for 60 minutes at room temperature. Following this fixation, a methanol dehydration/rehydration series was run (25%,50%,75%,95%,75%,50%,25%-10 minutes each). Samples were then incubated in PBS/.2% Triton X-100/1% BSA for 30 minutes followed by antibody addition in the same solution and incubation at 4°C overnight. The following day three 30 minute washes in PBS/.2% Triton X-100, with the first wash containing DAPI at 5 ug/ml, were carried out.

#### FITC-D-Alanine Labelling of Blautia coccoides

Following one day of co-culture on-chip, upper channel reservoirs were exchanged for new reservoirs containing 250µM FITC-conjugated D-alanine. Co-cultures continued for a second day, followed by chip disassembly from tubing. Chips were rinsed briefly with PBS and fixed in 2% paraformaldehyde for 15 minutes. Nuclei were stained with DAPI at a concentration of 1ug/ml.

The following antibodies were utilized: #3199 Cell Signaling Technology - E-Cadherin (24E10)(Alexa Fluor® 488) *1:100*, #4627 Cell Signaling Technology - Beta-Catenin (L54E2)(Alexa Fluor® 647) *1:100*, MA3-39100-A555 Invitrogen – ZO-1 (1A12)(Alexa Fluor™555) *1:100*, Santa Cruz Biotechnology sc-515106 AF488 – Mucin-2 (H-9) *1:50*, Abcam (ab206091) Alexa Fluor® 488 Cytokeratin-18 [EPR1626] *1:500*. Phalloidin Alexa Fluor 555 was used at .33 uM, DAPI at 1-5 ug/ml and D-Alanine-FITC at 250uM.

Antibody/Dye labelling images were collected on one of two imaging systems: 1) A Leica THUNDER widefield system on a Leica DMi8 inverted stand equipped with a 10x .32 NA HC PL Fluotar objective or 2) A Zeiss LSM 880 equipped with an AiryScan detector. Standard confocal mode was utilized for most images, however the AiryScan was utilized for imaging of *Blautia coccoides* labelled with D-Alainine-FITC in Figure 3L. Images on this system were acquired using either a 63x 1.4 NA oil immersion or 40x 1.2 NA water immersion objective.

### Chip Ab-/Adsorption Assay

Polycarbonate chips were prepared as stated above. Mock PDMS chips were prepared by casting Sylgard 184 at a ratio of 10:1 into molds which was baked for 4 hours. PDMS components were punched for fluid ports and bonded using plasma bonding. Each chip was loaded with 10µM rhodamine b and incubated at room temperature for two hours. Chips were rinsed with PBS, then washed four times for 10 minutes each in PBS. Images were gathered using a Leica Thunder widefield fluorescence microscope.

### Oxygen tension measurement

Gut chips cultured for six days were transitioned to white media. Prior to introduction, inlet media for upper channels was depleted of oxygen overnight in an anaerobic jar with Aneropack-Anaero anaerobic gas generator. Chips were cultured a further three days. On day 9 of chip culture, OXNANO probes (suspended in distilled water, then diluted in PBS and 10x PBS to maintain osmotic pressure) were introduced to the upper chamber at 100uL/min until probe solution reached collection syringes followed by flow at 100uL/hr for 10 minutes. Readings were taken at three points along the channel. Each reading lasted for two minutes taking readings every 5 seconds. Values at each point were averaged using PyroScience Data Inspector software. Readings were normalized in the software to the proportion of oxygen in ambient air, so output values were multiplied by 20.95% to report %oxygen in the solution. N=4, taken from two independent experiments.

## Acknowledgements

This project was supported by funding (Exploratory6781119, Full6950120 and Full 2021-1391) from the Cancer Early Detection Advanced Research Center at Oregon Health & Science University’s Knight Cancer Institute and the Collins Medical Trust in Portland, Oregon. The authors would like to acknowledge intellectual contributions by Katherine Huynh, Ravi Thakur, Parul Katoch, Beverly Emerson, and George Thomas, organoid culture assistance by Priyanka Singla and assistance of the staff in the OHSU Advanced Light Microscopy Core and the OHSU Knight Cancer Institute Advanced Multiscale Microscopy Shared Resource, supported by P30CA069533 (PI: Druker, B.). Figure schematics were generated using BioRender.com and Autodesk Fusion360.

## Supporting Information

Supporting information may be found online or by contacting the authors.

