## Supporting Information for "A Linkable, Polycarbonate Gut Microbiome-Distal Tumor Chip Platform for Interrogating Cancer Promoting Mechanisms"

**Supporting Methods**

*Extracellular Matrix (ECM) Functionalization*

ECM solutions including collagen from rat tail tendon at a concentration of 50µg/mL in either phosphate buffered saline (PBS) or 0.02M acetic acid or 50µg/mL each of collagen and fibronectin in 1%BSA were prepared and introduced to sterile polycarbonate membranes in chips. Coating was conducted at room temperature or in a tissue culture incubator for either 3 hours or overnight. Following coating, solution was aspirated and membranes allowed to dry for 30 minutes. For fibril deposition imaging, membranes were rinsed with PBS followed by imaging via two-photon microscopy. For cell seeding assays, conducted within a modified chip for static culture, dried membranes were incubated with media for one hour prior to application of cell suspension prepared in conditioned media. Cells were allowed to adhere for 2 hours, 4 hours, 6 hours, or overnight followed by a media wash and incubation prior to imaging. Cells were cultured on membranes for a total of two days followed by DAPI staining at a concentration of 1µg/mL for 30 minutes. Cells were rinsed twice with PBS then imaged via fluorescence microscopy and assessed for monolayer coverage.

*Organoid Culture On-Chip*

Organoids were prepared following protocols from the National Cancer Institute Patient-Derived Models Repository. Briefly, organoids were cultured in a complete media composed of Advanced DMEM/F12 with HEPES, GlutaMAX, Primocin, and L-WRN cell conditioned media further supplemented with N-acetylcysteine, nicotinamide, B-27, N-2 and Y-27632 dihydrochloride. To passage organoids, a simplified wash media (composed of advanced DMEM/F12, HEPES, GlutaMAX, Primocin and FBS) was supplemented with Dispase II and added to organoid cultures. Following incubation at 37$^{o}$C for 1.5-2 hours, cultures were dissociated by pipetting followed by neutralization in more simplified wash media and centrifugation. Supernatant was removed, wash media added and mixed with organoids, then centrifuged again. After removing supernatant, organoids were further dissociated by repeated pipetting then all media was removed. Remaining organoids were mixed with Reduced Growth Factor BME Type 2 on ice and injected into sterilized SCTC. Chips were incubated at 37$^{o}$C for 15 minutes to allow BME2 to set, followed by introduction of media and initiation of flow at a rate of 10.5uL/hr.

**Supporting Data**

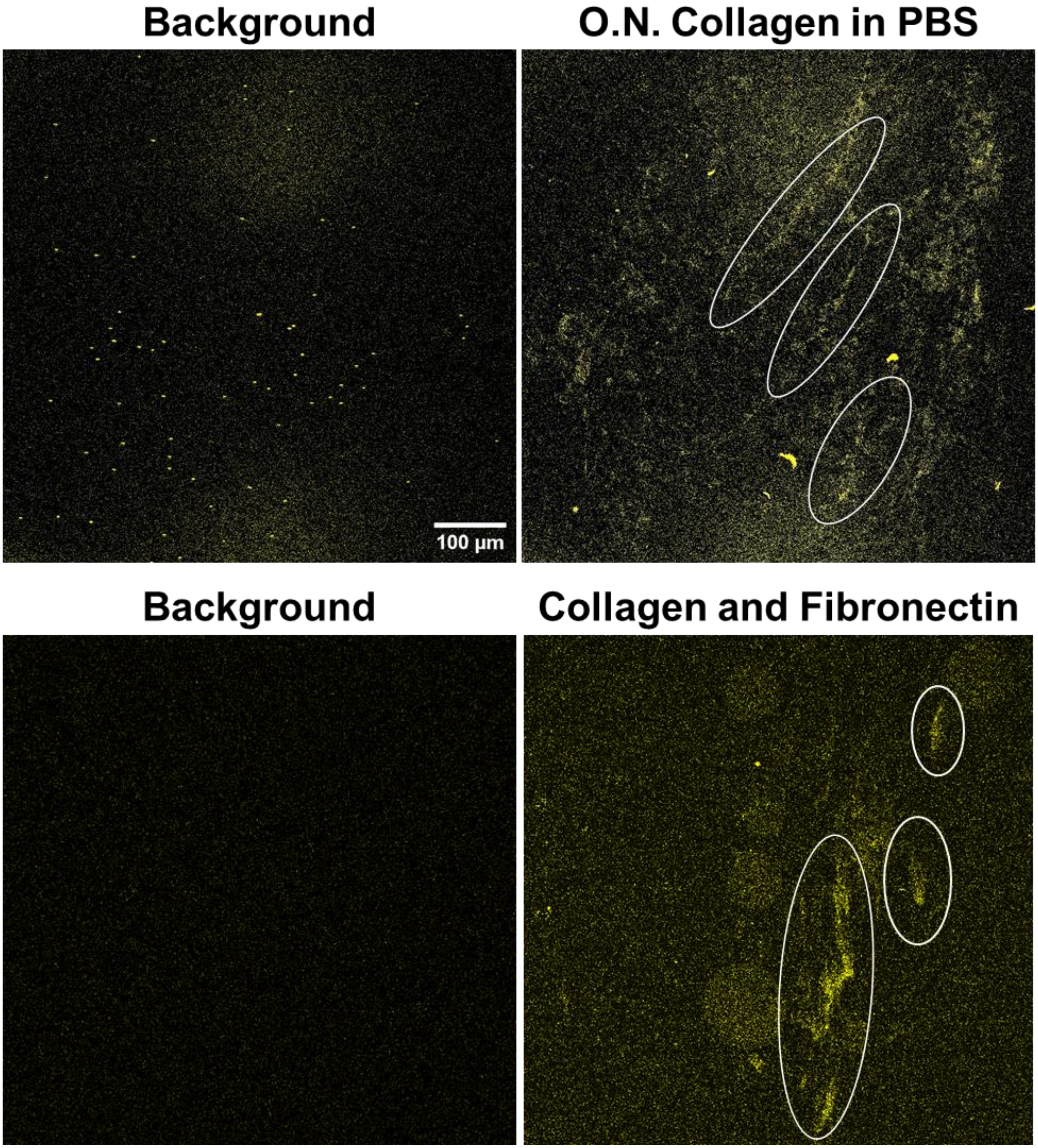


**Figure S1.** Imaging of collagen fibril deposition *via* two-photon microscopy. Examples of collagen fibrils are noted by white ovals.

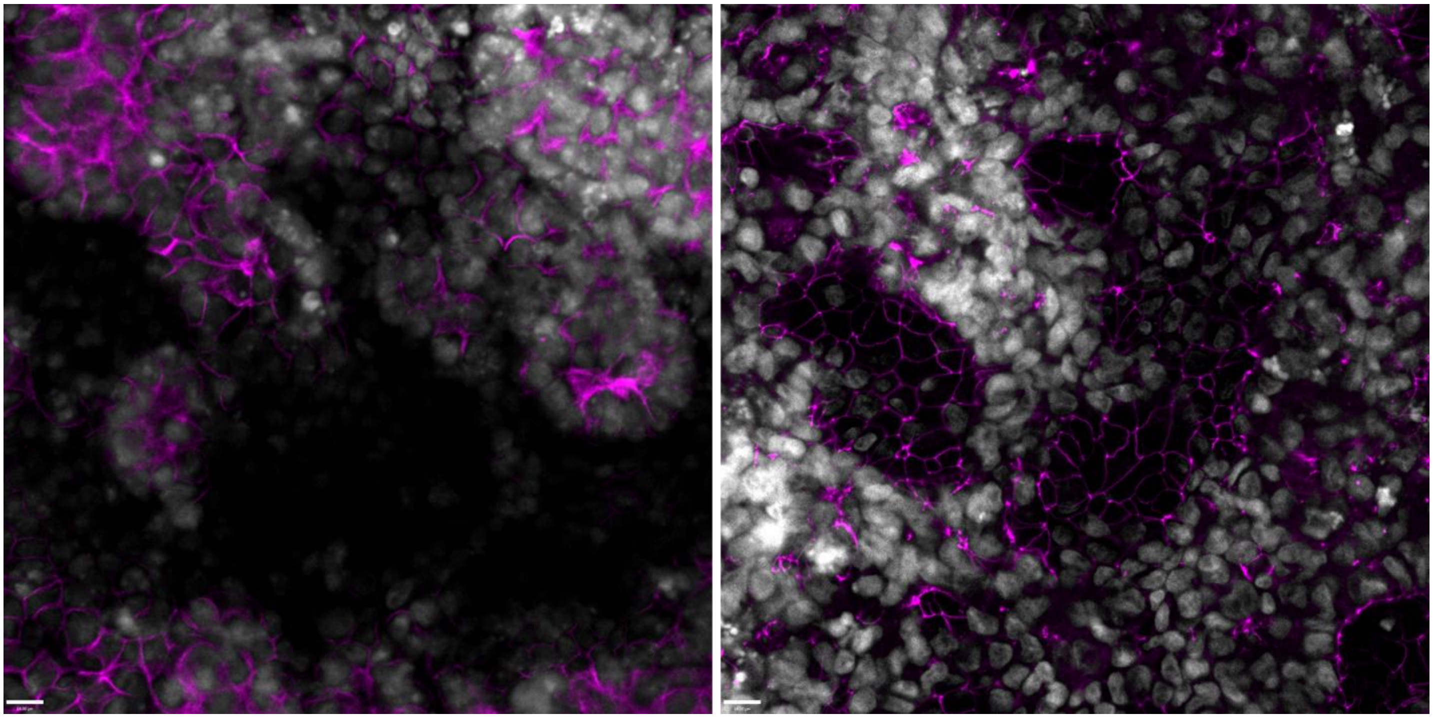

**Figure S2.** Confocal microscopy of gut chips stained for beta catenin(left) and ZO1 (right) showing a larger field of view of images in the text.

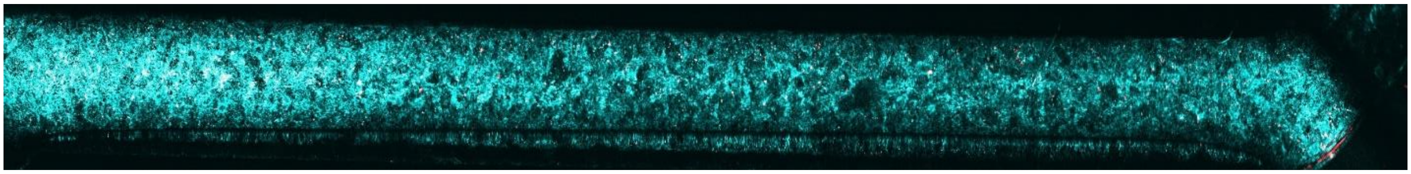

**Figure S3.** Two-photon microscopy image of gut epithelium after two days of co-culture with *Blautia coccoides*, staining for live cells (blue) and dead cells (red).

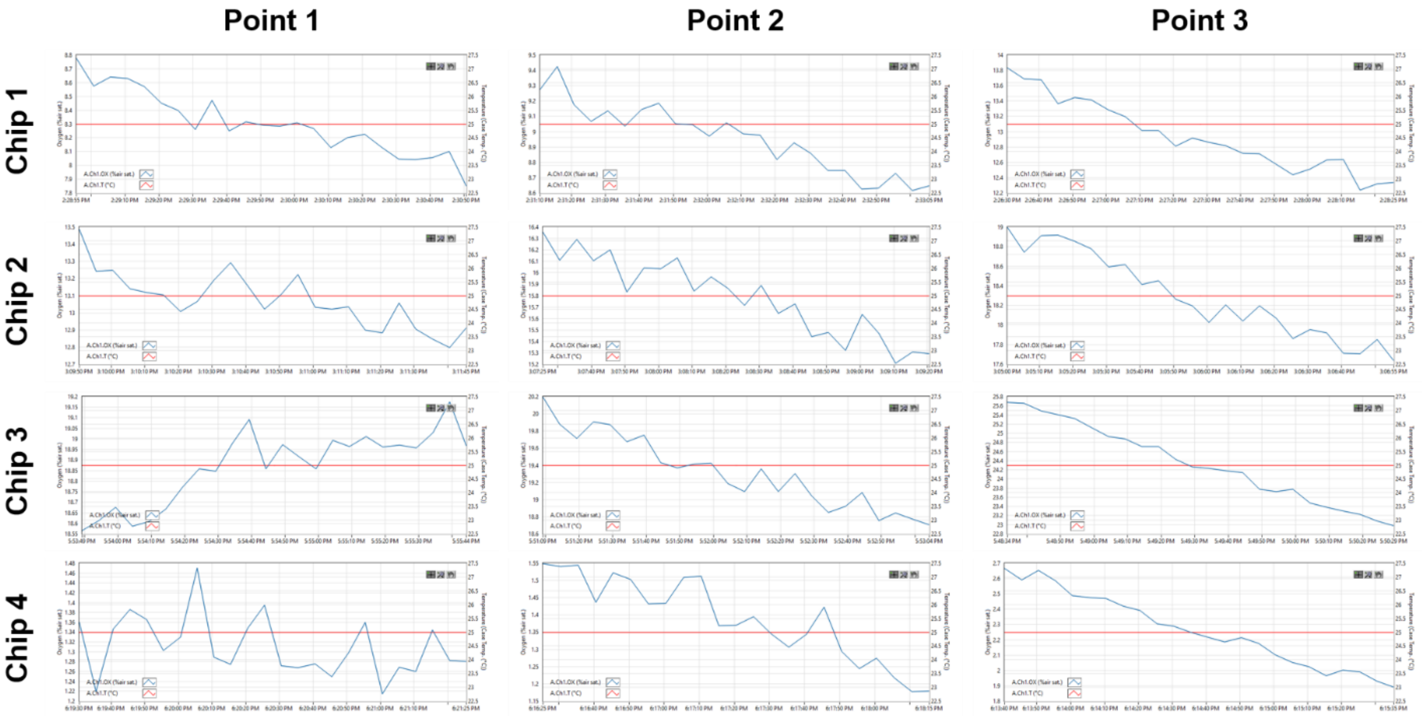


**Figure S4.** Readings taken from nine day-old gut chips using PyroScience's OXNANO bulk oxygen sensor flowing through the deoxygenated lumen chamber.

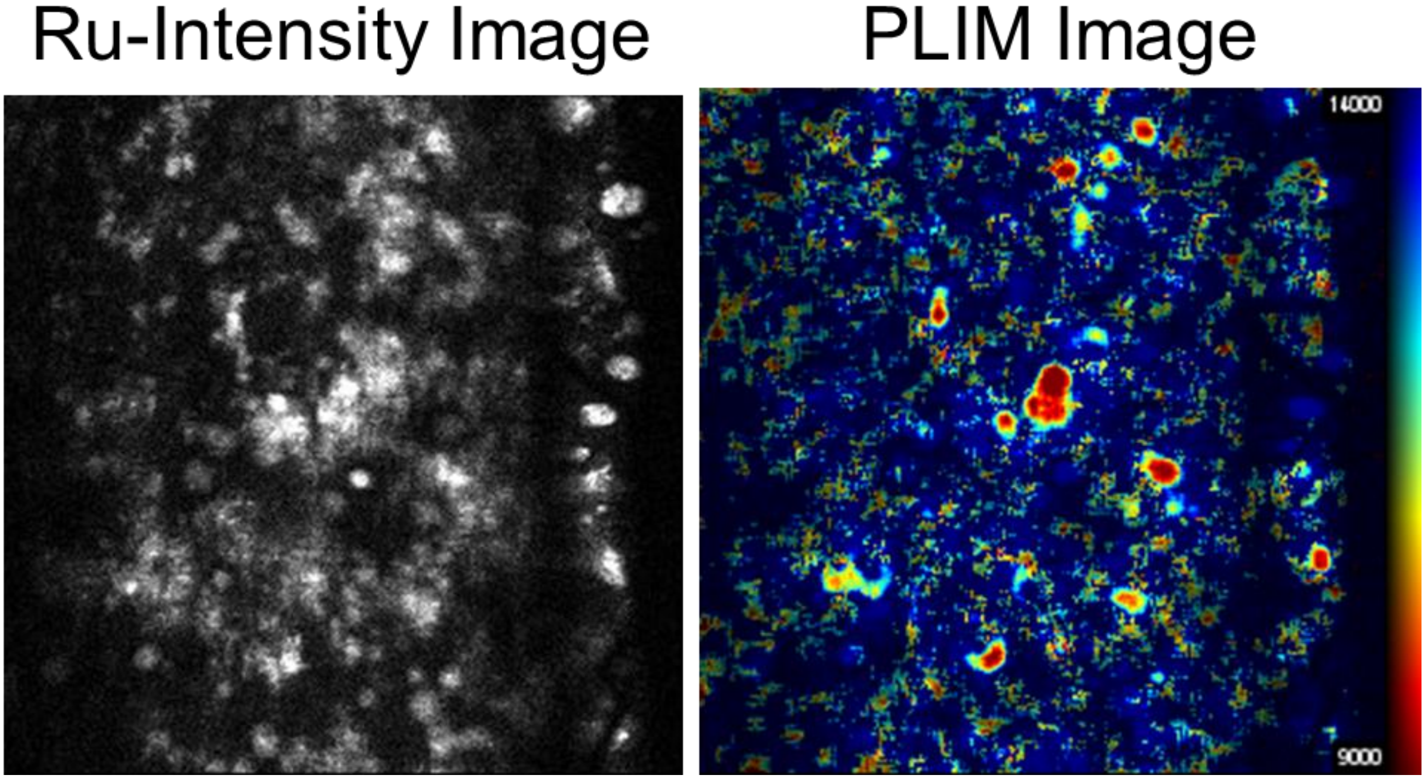


**Figure S5.** Phosphorescence lifetime imaging of oxygen sensors in a gut chip grown for ten days with three days of co-culture with *B. coccoides* shows oxygen gradients within the gut villus-like architecture.

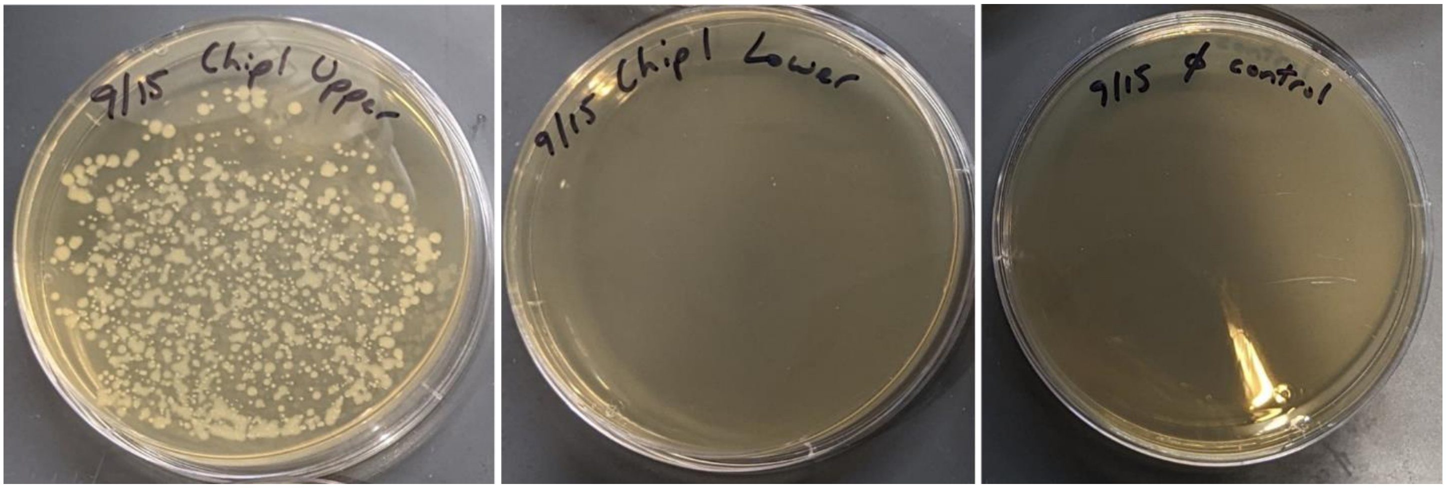


**Figure S6.** Plating media collected from the upper lumen compartment and the lower circulatory compartment of chips following two days of co-culture between Caco-2 epithelium with obligate anaerobe *B. coccoides* demonstrate bacteria maintain viability in the polycarbonate gut microbiome chip and do not extravasate into the circulatory compartment.

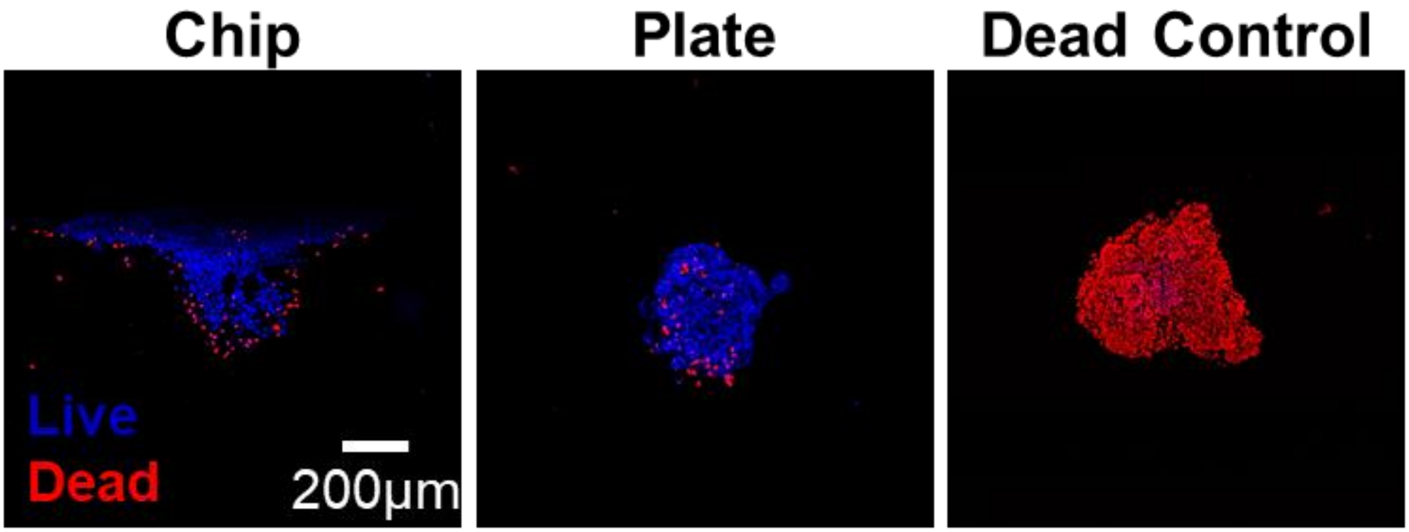


**Figure S7.** MCF7 spheroids cultured for one week in low adhesion u-bottom plates were loaded into SCTC with 1.45mg/mL collagen. Following culture at 60uL/hr flow for two days, three chips and three spheroids kept in plate culture were assessed for viability via staining with Hoechst for live cells (blue) and ethidium homodimer for dead cells (red). Chip culture did not reduce spheroid viability.

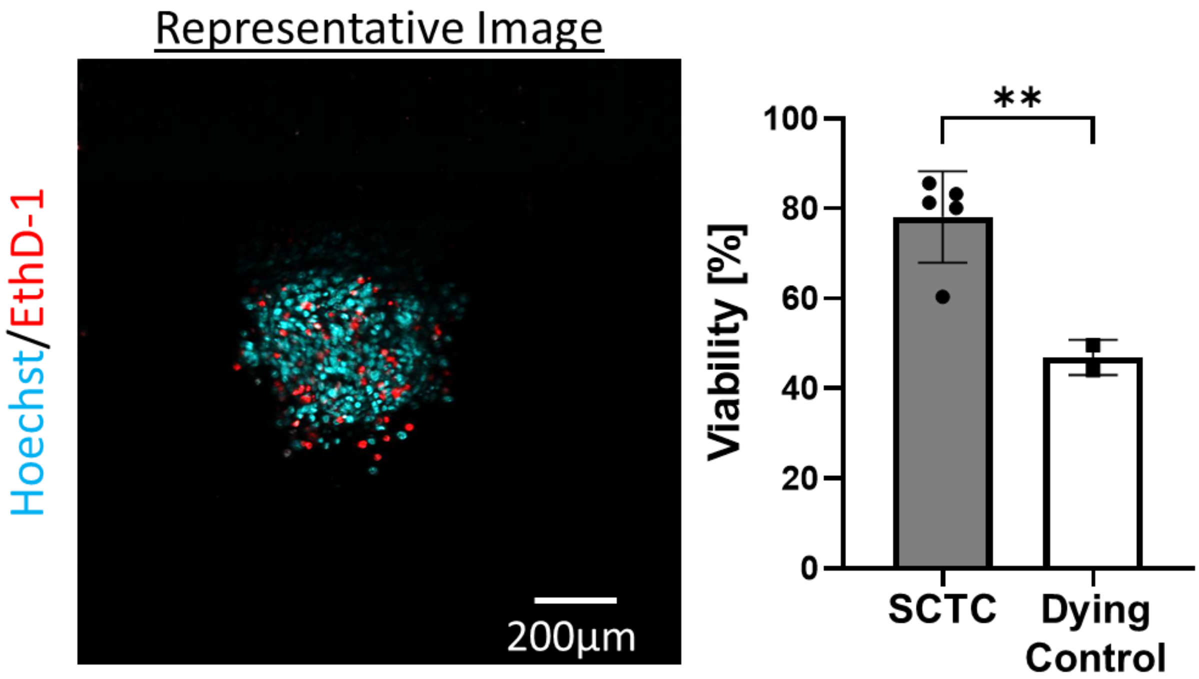


**Figure S8.** MCF7 spheroids cultured for five days in low adhesion u-bottom plates were loaded into SCTC with 2.7mg/mL collagen. Following culture at 100uL/hr flow for three days, three chips were assessed for viability with one dying spheroid control via staining with Hoechst for live cells (blue) and ethidium homodimer for dead cells (red). Spheroids maintained high viability when cultured on chip.

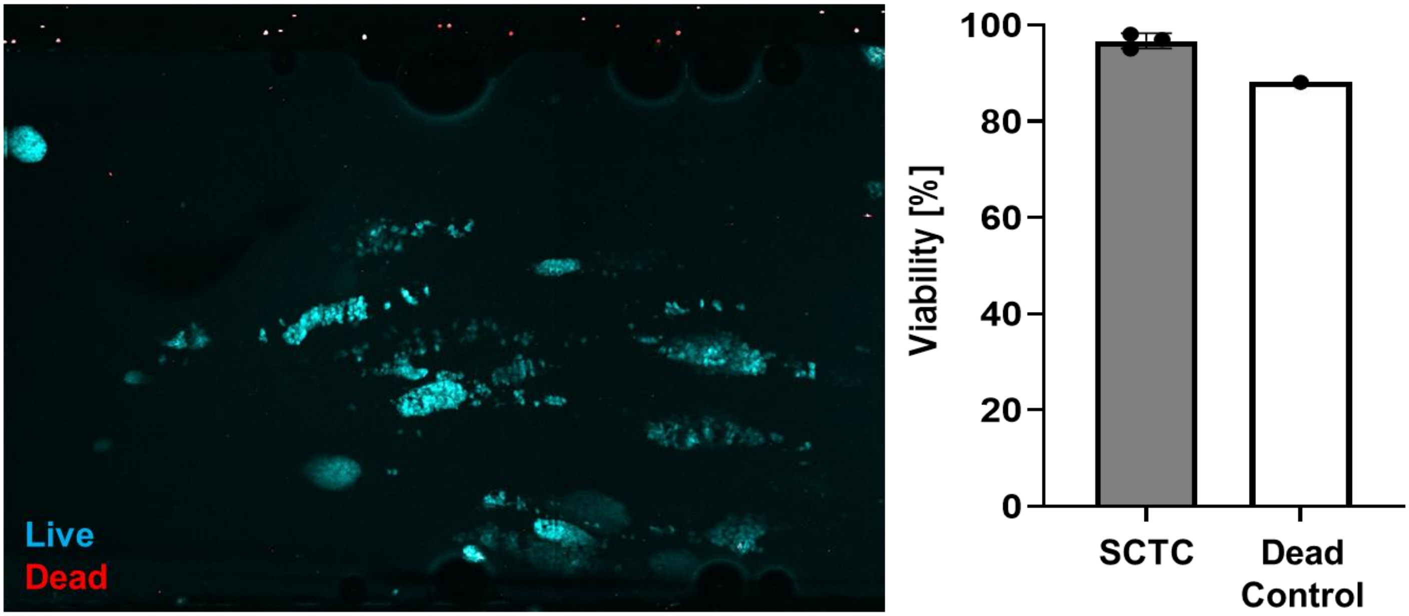


**Figure S9.** Kidney cancer patient-derived organoids were passaged and loaded into the SCTC in BME2. After letting hydrogel set, chips were cultured with constant flow at 10.5uL/hr for one week. Three chips were assessed for viability by staining with Hoechst for live cells (blue) and ethidium homodimer for dead cells (red). A fourth chip was treated as dying organoid control. Each chip value was determined by averaging values for multiple organoids. Organoids maintain high viability on-chip.

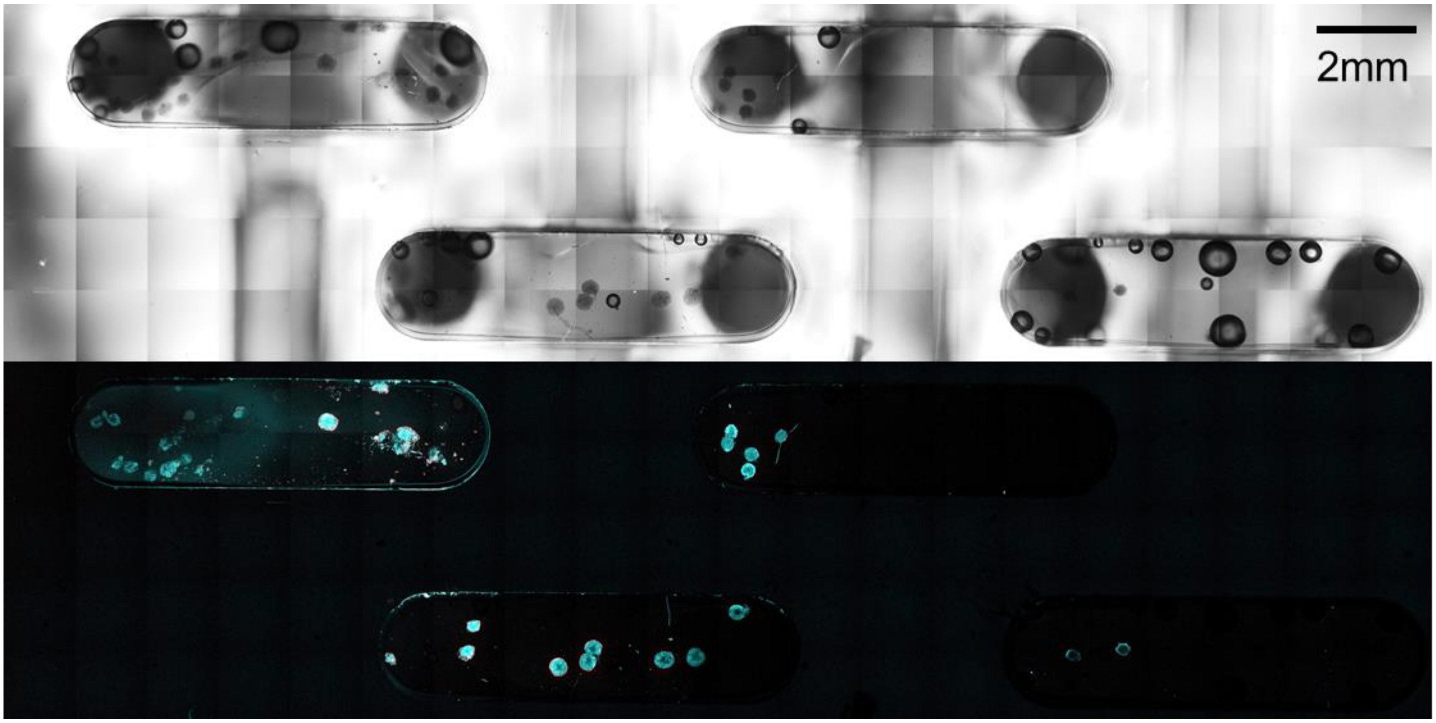


**Figure S10.** MCF7 spheroids cultured for five days in low adhesion u-bottom plates with transition to white media at day three were loaded into QuadTC in 2.7mg/mL collagen. Following culture under flow for two days, chip was assessed for viability via staining with Hoechst for live cells (blue) and ethidium homodimer for dead cells (red). Image shows stitched panels in brightfield (top) and fluorescence microscopy (bottom) to cover all four culture wells. Spheroids maintained high viability when cultured on-chip.

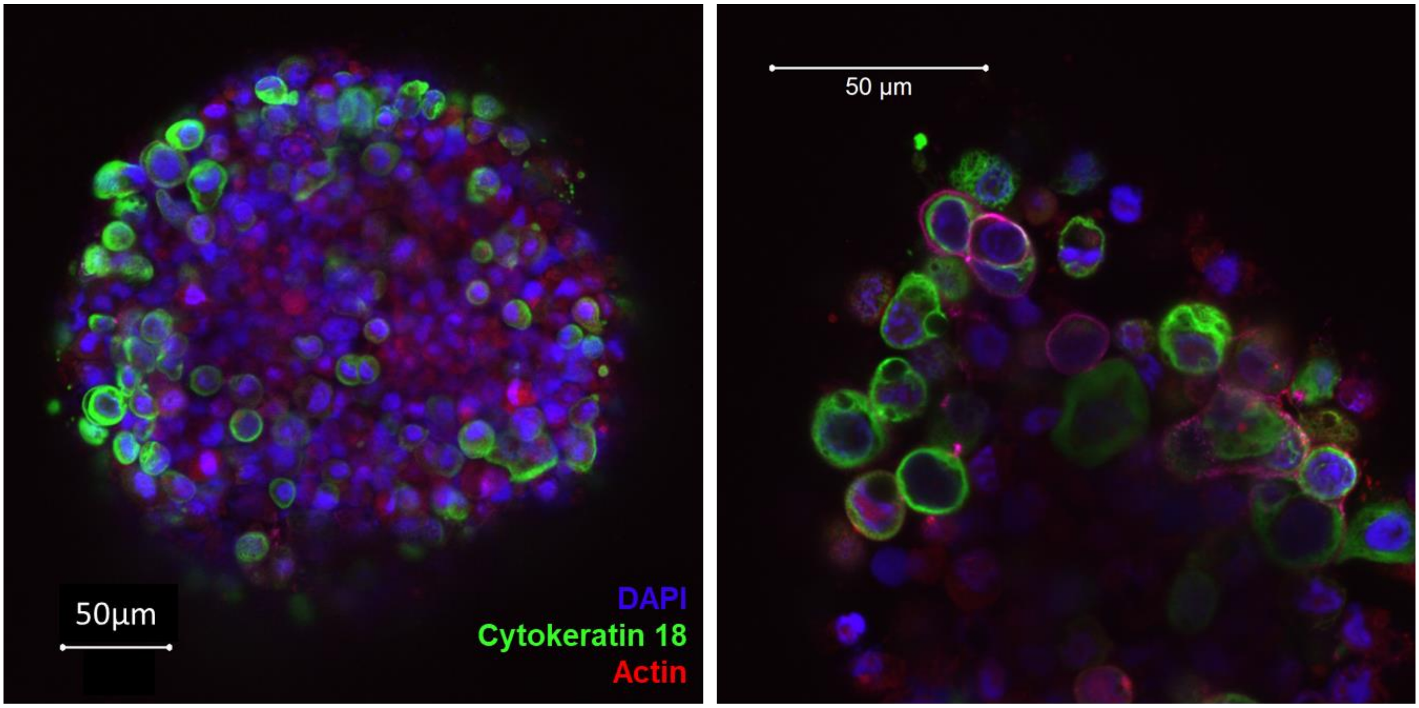


**Figure S11.** High resolution on-chip imaging in SCTC**.** MCF7 spheroids cultured for one week in low adhesion u-bottom plates were loaded into SCTC in 2.7mg/mL collagen. Following culture under flow at 60µL/hr for two days the chip was simultaneously fixed and permeabilized. Chip was dehydrated with methanol then rehydrated in PBS followed by blocking and staining using a tissue processing microwave. n=1

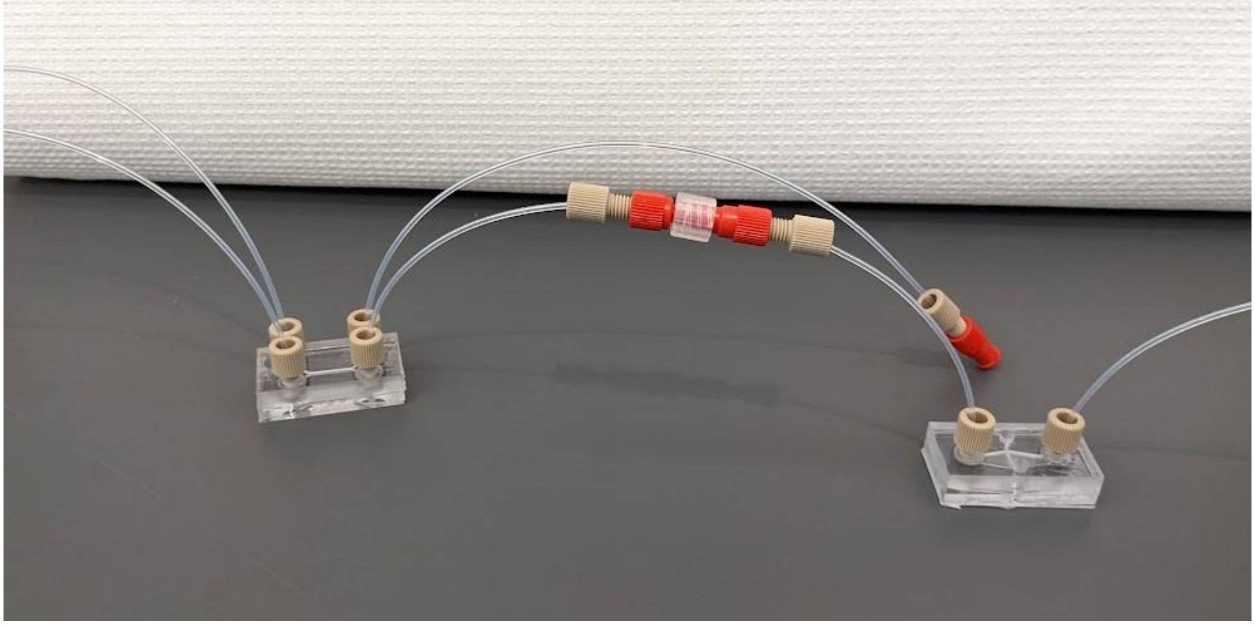

**Figure S12.** Example setup of gut microbiome and tumor chips linked in series for culture under flow driven by syringe pump.

**Video 1.** Gut chips culturing Caco-2 cells for one week were inoculated with *Blautia coccoides.* After one day of co-culture, inlet media was supplemented with 250µM FITC-D-alanine. Following another day of co-culture to allow incorporation of D-alanine into bacterial cell walls, chips were rinsed and fixed. Video depicts a z-stack of FITC-D-alanine labelled *B. coccoides* (green) growing at the surface of a Caco-2 villus-like structure, Caco-2 cell nuclei labelled with DAPI (blue).
